# Low centrosome numbers correlate with higher aggressivity in ovarian cancer

**DOI:** 10.1101/623983

**Authors:** Jean-Philippe Morretton, Anthony Simon, Aurélie Herbette, Jorge Barbazan, Carlos Pérez-González, Camille Cosson, Bassirou Mboup, Aurélien Latouche, Tatiana Popova, Yann Kieffer, Pierre Gestraud, Guillaume Bataillon, Véronique Becette, Didier Meseure, André Nicolas, Odette Mariani, Anne Vincent-Salomon, Marc-Henri Stern, Fatima Mechta-Grigoriou, Sergio Roman Roman, Danijela Matic Vignjevic, Roman Rouzier, Xavier Sastre-Garau, Oumou Goundiam, Renata Basto

## Abstract

Centrosome amplification, the presence of more than two centrosomes in a cell is a common feature of most human cancer cell lines. However, little is known about centrosome numbers of human cancers and whether amplification or other numerical aberrations are frequently present. To address this question, we have analyzed a large cohort of human epithelial ovarian cancers (EOCs) from 100 patients. Using state-of-the-art microscopy, we have determined the Centrosome-Nucleus Index (CNI) of each tumor. We found that EOCs show infrequent centrosome amplifications. Strikingly, the large majority of these tumors presented low CNIs. We show that low CNI tumors are enriched in the mesenchymal subgroup and correlate with poor patient survival. Our findings highlight a novel paradigm linking low centrosome number with highly aggressive behavior in ovarian cancers and show that the CNI signature may be used to stratify ovarian cancers.

## INTRODUCTION

The centrosome is the main microtubule-organizing center of animal cells. Each centrosome is composed of two centrioles surrounded by pericentriolar material (PCM), which is the site of microtubules nucleation. The centrosome facilitates the accuracy of chromosome segregation during mitosis and influences cell polarity and migration (Bettencourt-Dias and Glover, 2007; Bornens, 2012). Centrosome duplication is normally tightly controlled so that each centrosome duplicates only once per cell cycle (Bornens, 2012; Gönczy, 2015; Nigg and Holland, 2018). The presence of more than two centrosomes in a cell, centrosome amplification, is associated with tumorigenesis. T. Boveri proposed for the first time, more than one hundred years ago a link between extra centrosomes, multipolar divisions, and aneuploidy (Boveri, 2008). When induced by manipulating the centrosome duplication machinery, centrosome amplification is sufficient to drive tumor formation *in vivo* in various tissues in different animal models (Basto et al., 2008a; Coelho et al., 2015; Serçin et al., 2016; Levine et al., 2017).

Although centrosome amplification is generally associated with abnormal cell division and so aneuploidy (Boveri, 2008; Sabino et al., 2015; Serçin et al., 2016; Levine et al., 2017; Raff and Basto, 2017), centrosome amplification can also impact cellular homeostasis in alternative ways. For example, when induced in breast epithelial cells, centrosome amplification leads to the assembly of Rac1-dependent invasive protrusions (Godinho et al., 2014). Centrosome amplification can also drive cancer cell invasion in a non-cell-autonomous manner through increased oxidative stress (Arnandis et al., 2018). Non-cell autonomous detachment of mitotic tumor cells is described in organoids containing increased levels of Ninein-like protein, which induces centrosome structural defects (Casenghi et al., 2003; Schnerch and Nigg, 2016; Ganier et al., 2018). Even though numerical centrosome defects are described in different cultured cancer cell types (Marteil et al., 2018), very few studies have described centrosome number alterations in tumors *in situ* (Goundiam and Basto, 2021; Zyss and Gergely, 2009).

Epithelial ovarian cancers (EOCs) are the most lethal gynecologic malignancies (Berns and Bowtell, 2012). The high mortality rate is a result of late diagnosis and limited therapeutic options despite the use of new drugs, such as inhibitors of angiogenesis or DNA repair pathways (Konstantinopoulos et al., 2015; Pujade-Lauraine et al., 2017). 75% of EOC patients are diagnosed at advanced disease stages, resulting in low 5-year overall survival rate (Vaughan et al., 2011; Torre et al., 2018). The histological classification includes mainly serous, endometrioid, mucinous, and clear cells carcinomas. The most common EOCs subtype is high-grade serous (HGSOC), which presents a worse overall prognosis (Ramalingam, 2016). Moreover, up to 50% of HGSOC exhibit defects in homologous recombination (HR) pathways (Bell et al., 2011). HR deficient (HRD) patients with germline or somatic mutations in *BRCA1/2* genes are known to be more sensitive to platinum-based chemotherapy and Parp inhibitors than non-BRCA mutated tumors (Konstantinopoulos et al., 2015; Konstantinopoulos and Matulonis, 2018), more broadly defined as HR proficient (HRP) patients. In HGSOCs, four major transcriptomic signatures have been identified by TCGA consortium: Proliferative, immunoreactive, differentiated and mesenchymal. The mesenchymal subtype, also named stromal, angiogenic or fibrosis by other studies (Mateescu et al., 2011; Bentink et al., 2012; Verhaak et al., 2012; Huang et al., 2007; Kieffer et al., 2020; Integrated genomic analyses of ovarian carcinoma, 2011) is characterized by increased expression of Hox genes and stromal components. This group regroups HGSOCs with worse prognosis.

With the aim of characterizing the centrosome status in human cancers, we used a large EOCs cohort composed of 100 naive tumors comprising 88 HGSOCs. We used immunofluorescence and state-of-the-art microscopy to detect and quantify centrosome numbers. For each tumor, we established the centrosome-nucleus index (CNI) as a proxy to compare numbers among our cohort. Surprisingly, we found that the frequency of centrosome amplification was less important than could be predicted from the literature mostly based on cell culture. Additionally, we found that most tumors in our cohort contained cells without centrosomes, explaining the high frequency of low CNI tumors. Combining CNI data with genomic and clinical data revealed a striking association between low CNI and decreased patient survival.

## RESULTS

### Characterization of centrosome defects in human epithelial ovarian cancer (EOC) tissues

To analyze centrosomes in human epithelial ovarian cancers (EOCs), we obtained 20μm frozen tissue sections from the pathology department of Institut Curie. These were categorized as healthy tissues (corresponding to healthy ovaries from prophylactic oophorectomy or hysterectomy) or tumor tissues, including a mix of serous (90%), endometrioid (3%), mucinous (4%), and clear cell carcinoma (3%) (methods and Supplementary Table 1). All tumors were treatment-naïve, obtained after surgery without previous neo-adjuvant chemotherapy. Tissues were immunostained for Pericentrin (PCNT) and CDK5RAP2. These are two PCM components, and through their co-localization we can unambiguously identify centrosomes as defined in previous studies (Basto et al., 2008b; Serçin et al., 2016; Gambarotto et al., 2019). Using confocal microscopy, we obtained optical Z sections from ten random fields in the entire tissue (Figure 1A). Analysis of healthy tissues allowed us to identify centrosomes (Figure 1B). We also noticed the presence of structures that only contained one of the two centrosome markers (Figure 1B). These were not considered centrosomes. To further characterize and confirm the centrosomal configurations described above, we used 3D structural illumination microscopy (3D-SIM) of ovarian tissues labelled with the centriolar marker-Cep135 and PCNT, allowing higher resolution for both centrioles and PCM (Figure 1C). We found that in healthy tissues, each centrosome contained two centrioles and as expected (Conduit et al., 2015), PCNT surrounded one of the two centrioles, presumably the mother centriole.

**Figure 1.**
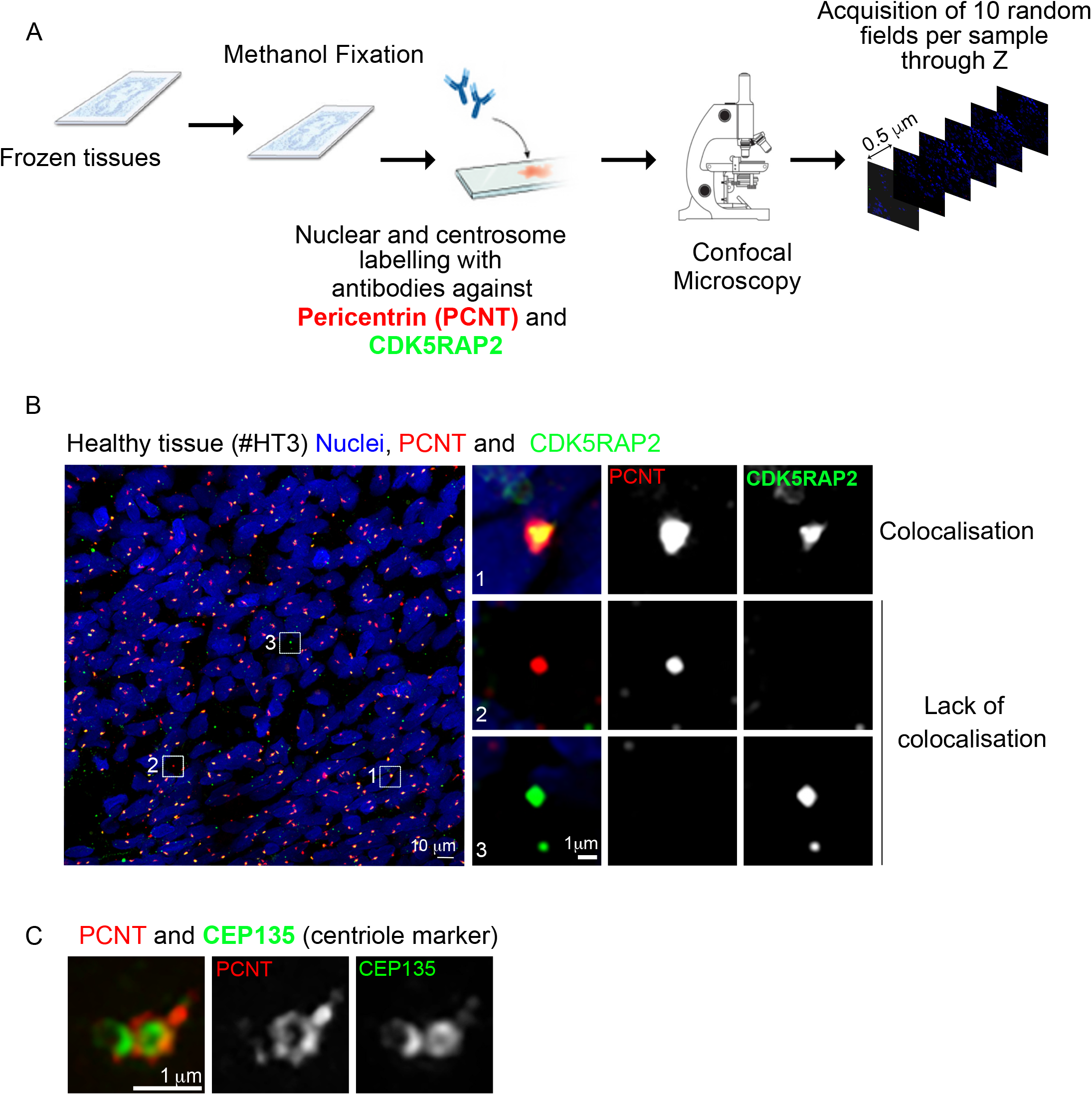
Characterization of centrosome numbers in healthy ovarian tissues. (A) Schematic diagram of the workflow used to analyze ovarian tissue sections. Frozen healthy or ovarian tumor tissues were sectioned into 20μm thickness sections and methanol fixed. These were subsequently immunostained for two centrosomes markers, and nuclei were labelled with DAPI. Ten random fields were imaged through the entire Z-stack using confocal microscopy. Each field was analyzed and centrosomes and nuclei were quantified manually. (B-C) On the left, representative image of low magnification view of healthy tissue immunostained with antibodies against pericentrin (PCNT) and CDK5RAP2, shown in red and green, respectively. DNA is in blue. Scale bar 10μm. The white dashed squares represent the regions shown in higher magnifications on the right. One centrosome was considered as such when PCNT and CDK5RAP2 signals co-localized. Lack of co-localization was noticed and discarded during quantification. (C) Super-resolution microscopy of healthy tissues immunostained for the centriole marker Cep135 (in red) and PCNT (in green). Scale bar 1μm.

Analysis of tumor tissues revealed the presence of highly heterogeneous phenotypes in respect to centrosome numbers and even overall aspect of the tissue (Figure 2A). This supports the requirement for the acquisition of multiple fields for each tumor. In most tumor sections, one or two centrosomes were readily noticed (Figure 2B top left insets). Surprisingly, however, in other nuclei, we could not detect centrosomes or even any signal from individual centrosome proteins (Figure 2B low left inset). In addition to the presence of few nuclei without centrosomes in certain sections, we noticed large regions containing nuclei without centrosomes (Figure 2C-left). In others tumor sections, these were restricted to a group of nuclei (Figure C-right).

**Figure 2.**
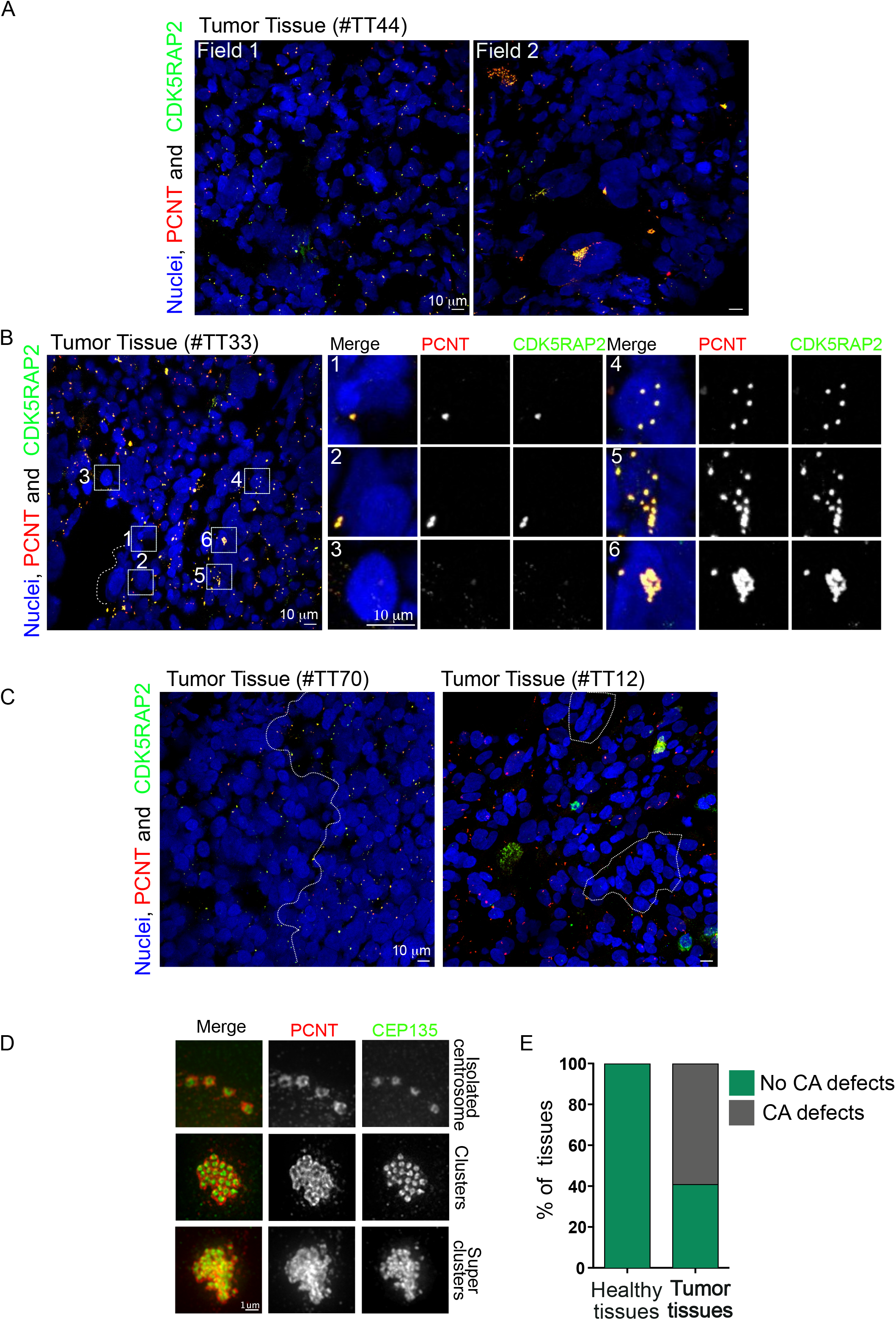
Characterization of centrosomes in EOCs. (A) Representative images of two different fields of the same tumor at low magnification immunostained with antibodies against pericentrin (PCNT) and CDK5RAP2, shown in red and green, respectively, DNA in blue, showing a high variability in nuclear size and centrosome number and size. Scale bar 10μm. (B) On the left, low magnification representative image of a tumor section labelled as described above. Scale bar 10μm. The white dashed squares represent the regions shown in higher magnifications on the right. (C) as in (B) to visualize large (left) and small (right) regions without centrosomes. Scale bar 10μm. (D) Super-resolution microscopy of tumor tissues immunostained for the centriole marker Cep135 (in red) and PCNT (in green). Scale bar 1μm. (E) Graph showing the quantification of the percentage of tissues with and without centrosome amplification (CA) of any type-Isolated centrosomes, clusters and super-clusters.

Considering centrosome amplification, in certain cells, extra centrosomes could be seen as isolated structures spread away from each other (Figure 2B, top right panel), and these were named isolated centrosomes. In other cells, they were clustered together - clustered centrosomes (Figure 2B, middle right panel). Interestingly, we also observed a configuration where ECs were tightly clustered in a single structure - super-clusters (Figure 2B, lower right panel). SIM analysis of these tumors, with the markers described above demonstrates the unusual extra centrosome morphologies (Figure 2D). Comparison between healthy and tumor tissues revealed the absence of centrosome amplification in healthy tissues, while a large fraction of tumors - 40%-did not have any defect (Figure 2E).

Altogether, the methodology employed to analyze 100 ovarian tumors and the comparison with healthy ovarian tissues revealed the unexpected presence of cells without centrosome and the presence of super-clusters.

### EOCs show low levels of centrosome amplification and many nuclei are not associated with centrosomes

We next quantified the frequency of these defects in a cohort of 19 healthy tissues and 100 tumor tissues. We only imaged and analyzed regions corresponding exclusively to the tumor, excluding the stroma that surrounds the tumor. Tumor tissues appeared very disorganized and it was difficult to ascertain the number of centrosomes per cell as, in many cases, centrosomes were not closely associated with the nucleus. To unmistakably quantify the centrosome number and to compare all tumors and healthy tissues, we visually counted the number of nuclei and the number of centrosomes in each field. We determined the Centrosome Nuclei Index (CNI) by dividing the number of centrosomes by the number of nuclei (Figure 3A). The data we present, therefore, is the result of manual counting.

**Figure 3.**
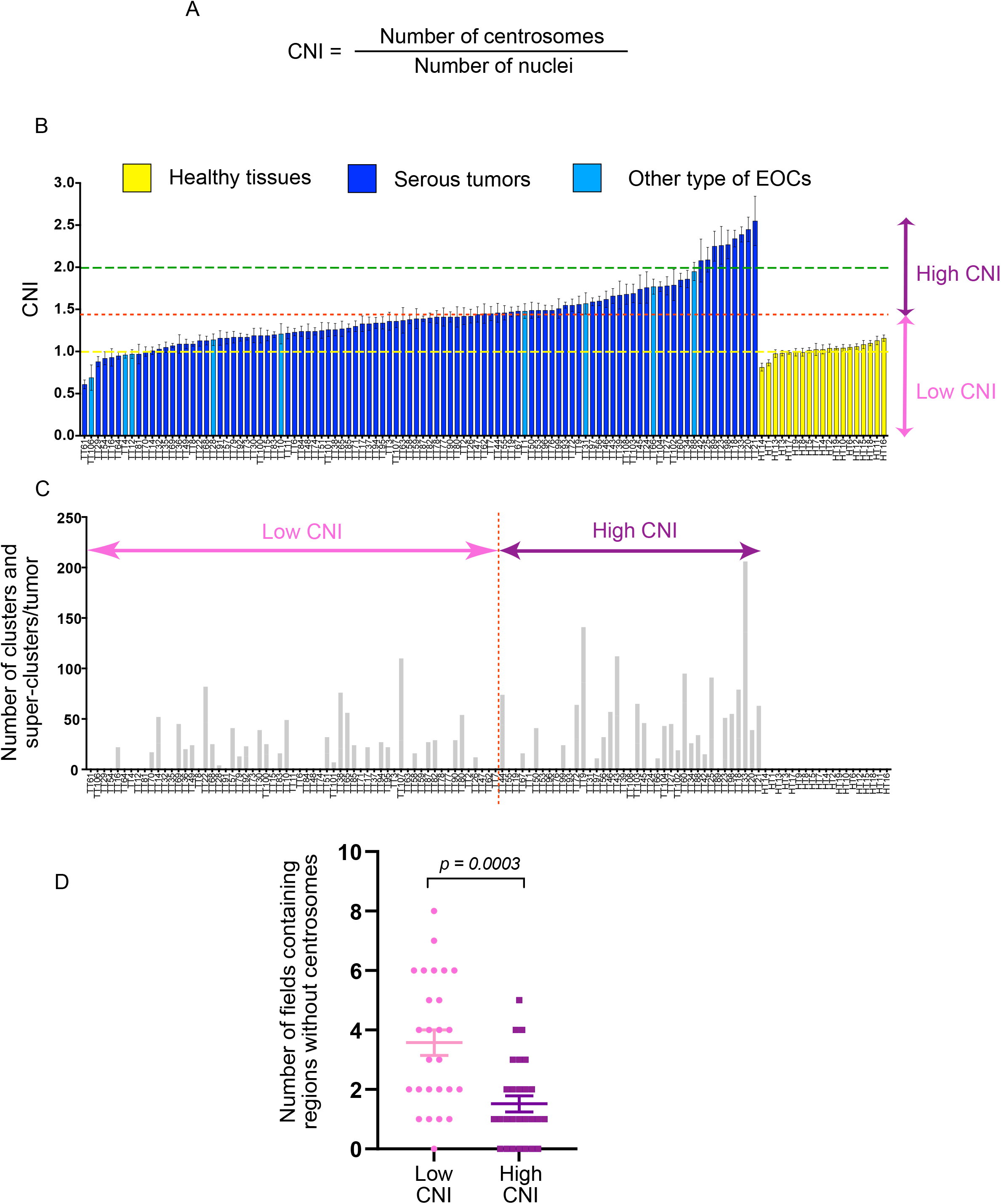
Characterization of the Centrosome Nuclei Index (CNI) in healthy and tumor tissues. (A) Diagram showing the CNI calculation. (B) Plot showing the CNI value of all 100 tumors (blue) positioned in ascending value and 19 healthy tissues (yellow) analyzed. The yellow dash line represents the average CNI value of all normal tissues analyzed (1.02) and the red line shows the threshold of CNI value defining HGSOCs as low or high CNI tumors (1.43). The green dash line represents centrosome amplification as defined by the literature (>2 centrosomes in the cell). (C) Plot showing the total number of clusters and super-clusters identified in the tumor population set. Note that the order of the tumors is conserved between the two plots to allow for comparison between the CNI and the number of extra centrosomes within the same tumor. (D) Dot plot graph showing the quantification of fields without centrosomes in high and low CNI tumors. Statistical significance was determined with a Mann-Whitney-test.

Overall, our analysis comprised 653627 nuclei, 874766 centrosomes from 1174 fields, with an average of 5248 nuclei counted per tumor. In healthy tissues, the average CNI was 1.02±0.02, and it was relatively stable, varying from 0.81 to 1.16 (Figure 3B). In tumors, however, the CNI was much more variable. On average, 1.43±0.04, with the minimum at 0.61 and maximum at 2.55. Interestingly, 89% of the tumors presented a CNI superior to the average CNI found in healthy tissues (Figure 3A, yellow dashed line, and Supplementary Figure 1A). However, only 9% of tumors exhibited extensive centrosome amplification with a CNI above 2 (Figure 3B, green dashed line), when defined by the presence of more than two centrosomes per cell (Boveri, 2008; Godinho et al., 2009; Marthiens et al., 2012).

We then restricted the subsequent analysis of our cohort to the high-grade serous ovarian cancers (HGSOCs). We dichotomized our population into two groups using Classification And Regression Trees (CART) method. This resulted in the categorization of the cohort into low CNI (≤ 1.45) and high CNI (> 1.45), with the majority of tumors −55 -falling into the low CNI category. 33 tumors were placed in the high CNI category (Figure 3B, red line). To evaluate the performance of the CNI as a classifier and to validate the 1.45 threshold, we used predictiveness curves (Huang et al., 2007). We performed this analysis considering the presence of extra centrosomes as 63% (n=56 out of 88) in the HGSOC cohort used in this study. We applied an unbiased resampling process considering early relapse, as this is a clinical parameter frequently used to characterize disease occurrence (Lheureux et al., 2019). The predictiveness curves showed that the optimum CNI value is 1.456 (95% CI= [1.22-1.76]), which confirms the threshold of 1.45 described above.

We first investigated whether the dichotomization of our tumor cohort in low and high CNI identified any preference for the different extra centrosome categories (isolated, cluster and super-cluster) identified by confocal microscopy. Using multivariate analysis, we recognized a significant trend for isolated centrosomes and clusters (p= 0.021 and p= 0.035 respectively, Supplementary Figure 1B) associated with high CNI tumors. However, even if not statistically significant, super-clusters tended to be associated with low CNI tumors (p=0.0788).

To gain more information about the distribution of clusters and super-clusters we plotted their number in parallel to the CNI analysis (Figure 3C). We found that certain tumors with low CNI (placed at the left side of the graph) contained clusters and super-clusters at similar frequencies as tumor tissues with high CNI (positioned at the right end of the graph). If both low and high CNI tumors can display similar numbers of clusters and super-clusters (which account for centrosome amplification), low CNI tumors must contain a higher frequency of cells without centrosomes. To further confirm this possibility, we randomly analyzed low and high CNI tumor fields and interrogated if these contained regions without centrosomes. Analysis of 10 fields from 26 low and 27 respectively high CNI tumors, revealed a higher frequency of areas presenting nuclei without any centrosome in low CNI tumors (Figure 3D).

Thus, EOCs are highly heterogeneous in terms of centrosome numbers. Surprisingly, only a small population of tumor cells display extra centrosomes and many nuclei lack centrosomes, which is unexpected in tumors of epithelial origin.

### CNI does not correlate with proliferation, mitotic index or genomic alterations in HGSOCs

We next explored the possible correlation between CNI and different molecular and clinical parameters. We analyzed whether the CNI status correlated with cell proliferation, using two indicators, the mitotic index (MI) and the proliferation marker Ki6. We did not find any correlation between CNI and MI or CNI and Ki67 signal (Supplementary Figure 2A-B).

Genomic alterations are frequently found in HGSOCs (Bell et al., 2011; Goundiam et al., 2015). Centrosome defects can lead to mitotic errors, chromosome instability and aneuploidy (Ganem et al., 2007; Pihan, 2013). To identify a possible link between centrosome number and genomic alterations, we used high-resolution Cytoscan arrays and Genome Alterations Prints (GAP) tools (Popova et al., 2009). We analyzed chromosome content (ploidy) and the presence of small and/or large DNA structural rearrangements. Importantly, we did not find any correlation between CNI status and ploidy, chromosome number, and DNA structural rearrangements (Supplementary Figure 2C-F).

Different pan-cancer studies (Zack et al., 2013; Bielski et al., 2018) have shown that whole-genome duplications (WGD) precede many types of genomic alterations. WGDs might represent a mechanism to generate aneuploidy, leading to chromosome number reduction, as shown in a mouse ovarian cancer model (Lv et al., 2012). WGD-positive (near tetraploid) tumors contain a ploidy of 3.31 on average, while ploidy is closer to ∼1.99 (near diploid) for WGD-negative tumors (Zack et al., 2013). We examined if CNI correlated with ploidy in our tumor cohort. We found that this was not the case (Supplementary Figure 2G), even though tumors with low CNI contained twice more near tetraploid (67%, n=26 out of 39 tumors) than near diploid karyotypes (33%, n=13 out of 39 tumors). In tumors with high CNI, however, the distribution was similar for near tetraploid tumors (46%, n=13 out 28 tumors) and near diploid tumors (54%, n=15 out of 28 tumors), (Supplementary Figure 2G).

Our analysis shows that MI and proliferation do not correlate with the number of centrosomes in EOCs. This is also the case for small or large chromosome breaks, suggesting a lack of correlation between the CNI and structural abnormalities. Interestingly, even if not statistically significant, low CNI seems to be associated with WGDs and hence with worse clinical prognosis (Bielski et al., 2018).

### High CNI correlates with better overall patient survival and high CNI tumors are frequently Homologous Recombination Deficient (HRD)

To further ascertain a correlation between CNI and patient outcome, we plotted HGSOC patient survival curves according to the CNI status. We found that low CNI was associated with worse overall survival (Figure 4A, Log-rank test: *p=0.018*, HR=1.931, 95% CI= [1.14-3.28]). Further, low CNI was also associated with a shorter relapse time after chemotherapy (Figure 4B, Log-rank test: *p=0.018*, HR=1.706, 95% CI= [1.059-2.750.28). In contrast, high CNI was associated with better overall survival.

**Figure 4.**
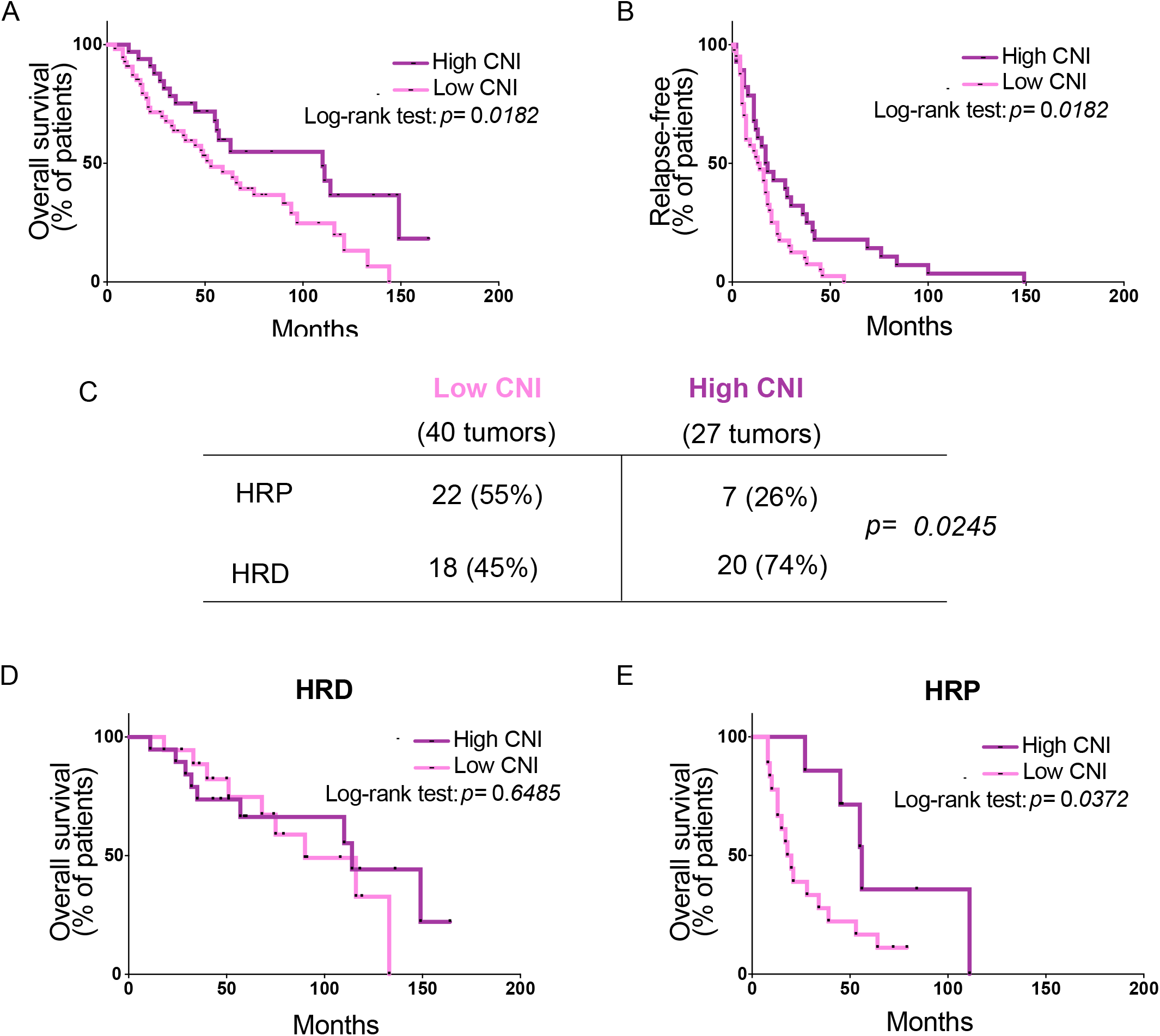
Lower CNI correlates with decreased survival and time to relapse. (A-B) Kaplan-Meier curves showing overall patient survival (A) and the percentage of patients without relapse, after the first line of chemotherapy (B) according to CNI status. Statistical significance was assessed with the Log-rank test for group comparison. (C) Contingency table showing the distribution of HR proficient (HRP) or deficient (HRD) in low and high CNI tumors. p-value from Fisher’s exact test. (D-E) Kaplan-Meier curves showing patient overall survival according to CNI status in HRD (D) and HRP (E) patients. Statistical significance was assessed with the Log-rank test.

To avoid any bias inherent to tumor stage, we investigated whether the CNI status reflected a particular stage taking into consideration the (International Federation of Gynecology and Obstetrics) FIGO classification (Prat, 2015). Importantly, we found that both low and high CNI tumors could be identified at all stages (I to IV) (Supplementary Figure 2H, Fisher test ns *p=0.07*). Interestingly, the majority of the cases in our cohort corresponded to stage III (59.0%, Supplementary Table 1), and these comprise low and high CNI tumors. We concluded that the association between high CNI and patient survival did not depend on tumor stage.

Mutations in genes encoding members of the DNA damage repair (DDR) pathway such as BRCA1 and BRCA2 (BRCA1/2), which are involved in homologous recombination (HR) lead to increased risk of breast and ovarian cancers (Chen and Parmigiani, 2007). We investigated an association between HRD and CNI status using the Large-scale transition (LST) genomic signature (Popova et al., 2009; Manié et al., 2016). This signature is based on the presence of large-scale chromosome breakpoints of at least 10Mb, which is an indicator of HRD. While low CNI tumors contained similar distributions of HRD and HR proficient (HRP) tumors (45% and 55% respectively), high CNI was mainly associated with HRD tumors (74% HRD and 26% HRP, respectively *p= 0.024*) (Figure 4C).

We next analyzed the overall survival of HRD patients. Notably, there was no significant association with the CNI status (Figure 4D, p=0.648, HR=1.229 and 95% CI= [0.49-3.17]). However, in HRP patients, who present a worse prognosis, significant differences according to CNI were noticed (Figure 4E, p=0.0372, HR=2.644 and 95% CI= [1.11-6.2]).

Overall and unexpectedly, these results show that an increased centrosome number within a tumor can be of better prognosis for HRP patients. Further, while the CNI status does not seem to be a parameter necessary to consider in HRD patient survival (for those who respond better to treatment), it can differentiate less sensitive HRP patients. These results suggest that the CNI index can be used to stratify HRP patients.

### Low CNI ovarian cancer spheroids do not show increased invasion or migration capacity

The lack of correlation between low CNI in tumors and mitotic index or chromosome abnormalities prompted us to explore other cellular mechanisms that may be influenced by low centrosome numbers. We decided to perform these experiments *in vitro,* in an isogenic background and using ovarian cancer cell lines. We generated inducible-(i) OVCAR8-PLK4 and SKOV3-PLK4 stable cell lines, where the expression of PLK4, the master centriole duplication, can be modulated by drugs. To increase centrosome numbers, PLK4 over-expression (PLK4OE) can be induced using doxycycline (Dox), as shown previously (Holland et al., 2012). To decrease centrosome number and thus conditions that mimic low CNI in a population of cells, we used centrinone, a PLK4 inhibitor (Wong et al., 2015). These cells will be referred to as centrinone cells. Treatment of either cell line with Dox or centrinone effectively impacted the CNI (Supplementary Figure 3A-D). Although proliferation was decreased in Dox and centrinone treated cells, these cells still proliferated (Supplementary Figure 3E), and apoptosis was only mildly increased (Supplementary Figure 3F-G). OVCAR8 and SKOV3 are EOCs cell lines with mutations in p53, explaining the continued proliferation in response to centrosome number alterations, in contrast to diploid untransformed cell lines (Holland et al., 2012; Lambrus et al., 2015; Wong et al., 2015).

It has been shown that centrosome amplification induces invasive features in a 3D culture mammary cell (MCF10A) model, both in a cell-autonomous and non-cell autonomous manner (Godinho et al., 2014; Arnandis et al., 2018). These cells show increased levels of an activated form of the small GTPase-RAC1 (Godinho et al., 2014). In EOC cell lines, however, we did not observe any significant difference in the levels of activated RAC1 after Dox or centrinone treatments (Supplementary Figure 4A-D).

EOCs undergo a particular mode of dissemination. Tumor cells detach from the primary tumor site, adhere to and migrate through the mesothelial cell layer that encloses peritoneal organs (Kipps et al., 2013) resulting in peritoneal metastasis (Iwanicki et al., 2011; Barbolina, 2018). Since we found that low CNI is a frequent characteristic of EOCs, we investigated if centrosome loss influenced mesothelial cell clearance. We performed these experiments using two different cell lines-iOVCAR8 and iSKOV3 treated with centrinone to induce low CNI (Supplementary Figure 3C-D and Supplementary Figure 5A-B) and used DMSO as a control (Supplementary Figure 3B-D). Both cell lines were grown as 3D spheroids and plated on top of mesothelial cells (Figure 5A). Time-lapse imaging allowed us to record the behavior of cancer cell spheroids over time for a period of 12hrs. Larger spheroids cleared mesothelial cells more rapidly and so we normalized clearance considering as the ratio between the final aperture and initial spheroid size (Figure 5B-C). Although iOVCAR8 and iSKOV3 cells cleared at different rates, decreased centrosome numbers did not influence mesothelial cell clearance when compared to controls (Figure 5D and Supplementary Figure 5A-B).

**Figure 5.**
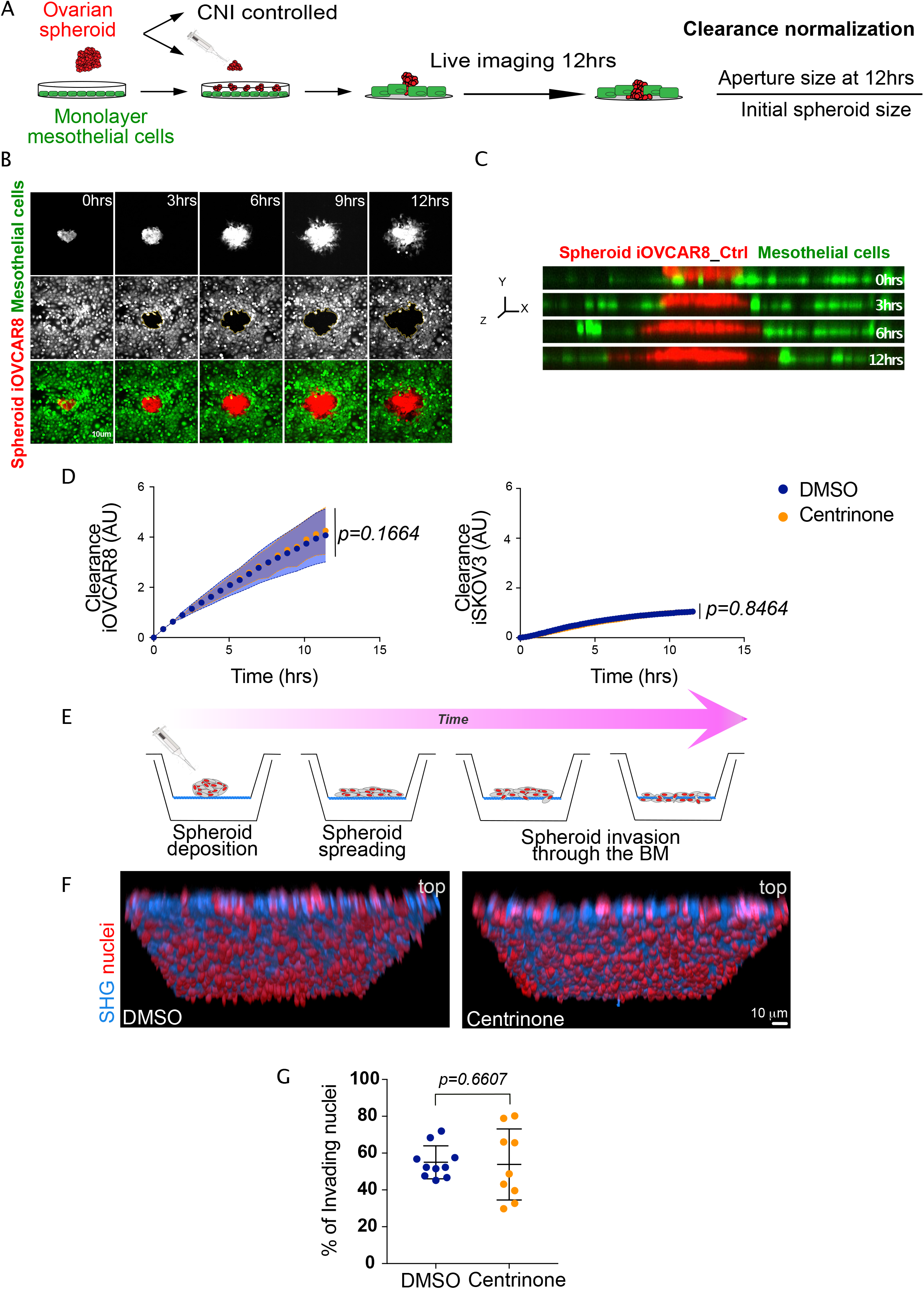
Ovarian cancer cells with low CNI do not show increased capacity to clear the mesothelium or invade the basement membrane. (A) Schematic diagram of the workflow. iOVCAR8 and iSKOV3 cancer cell lines were grown on polyHEMA to form spheroids and were labelled in red. Mesothelial cells were labelled in green and plated as monolayers on collagen I coated surfaces. After spheroid deposition on top of mesothelial cells, time-lapse movies of 12hrs were performed. The normalized clearance quantification was determined by dividing the hole size by the initial spheroid size at different time points. (B) Stills of a time-lapse movie of Ctrl spheroids. Time is shown in hours (hrs). Scale bar 10μm. (C) z-view of Ctrl cells as shown in B. Note the red-colored cancer cells at the beginning of the movie on top of the mesothelial layer while at later time points, they have cleared through the mesothelial cells. (D) Graph bars of the normalized clearance in A.U. of iOVCAR8 (top) and iSKOV3 (bottom) spheroids after the indicated treatments. For each experimental condition, at least 45 different spheroids were analyzed from three independent experiments. Statistical significance was assessed with the Anova test. (E) Schematic diagram of the experimental setup used in the basement membrane invasion assays. iSKOV3 cancer cell lines were grown on polyHEMA to form spheroids that were plated on top of basement membrane (BM) chambers prepared from female mice. Invasion was determined using Imaris BitPlane software. (F) Representative images of DMSO (left) and centrinone (right) spheroids. Red nuclei represent false colored invading nuclei. Scale bar 10μm. (G) Dot plot showing the quantification of the number of nuclei detected on the bottom side of the BM. For each experimental condition, three positions of three basement membrane inserts were analyzed. Statistical significance was assessed with the Mann-Whitney test. For all the experiments, the CNI was verified in parallel to confirm the low CNI conditions compared to DMSO.

Next, we asked whether low CNI influences basement membrane invasion. To address this question, we used decellularized mouse mesentery as an *ex-vivo* model, that replicates the complex basement membrane (BM) architecture located beneath the mesothelium (Glentis et al., 2018; Schoumacher et al., 2013) (Figure 5E). Cancer cell spheroids were plated on mesenteries and after seven days, we quantified invasion by counting the number of cells on the other side of the mesentery. We found that low CNI spheroids have the same invasion capacity as control spheroids (Figure 5 F-G and Supplementary Figure 5C).

Overall, our results show that centrosome loss does not impact migration or invasion in the ovarian cell models used in this study.

### Low CNI tumors correlate with mesenchymal tumor subtype

Our results so far showed that low CNI tumors correlate with worse prognosis as they present decreased overall survival and shorter relapse time after chemotherapy (Figure 4A-B). Molecular signatures based on gene expression data have defined different HGSOC molecular subtypes (Bell et al., 2011; Weinstein et al., 2013). Different signatures have also been identified in the cohort of patients used in this study, named stress and fibrosis signatures, which correspond to the expression of oxidative stress genes and mesenchymal genes, respectively (Mateescu et al., 2011; Weinstein et al., 2013). The fibrosis/mesenchymal subtype correlates with poor prognosis.

Since low CNI tumors are of worse prognosis, we investigated if there is an enrichment for mesenchymal signatures. This tends to be the case as, even if both low and high CNI tumors have a higher percentage of tumors expressing mesenchymal signatures, they tended to be enriched in low CNI tumors (Figure 6A). In our cohort of 72 HGSOCs with molecular characterization available, 45 tumors were low CNI (Figure 6B). Almost one third of these (27%, 12/45) were classified as mesenchymal according to the overlap signatures defined by (Mateescu et al., 2011; Bell et al., 2011) ( (Figure 6B-C). The number of high CNI tumors displaying the same signature was inferior (15%, 4/27), suggesting that more low CNI tumors overlap with the mensenchymal subtype. To challenge this approach, we interrogated the CNI status of all the mesenchymal tumors from our cohort. Importantly, 75% of these showed a low CNI (Figure 6D).

**Figure 6.**
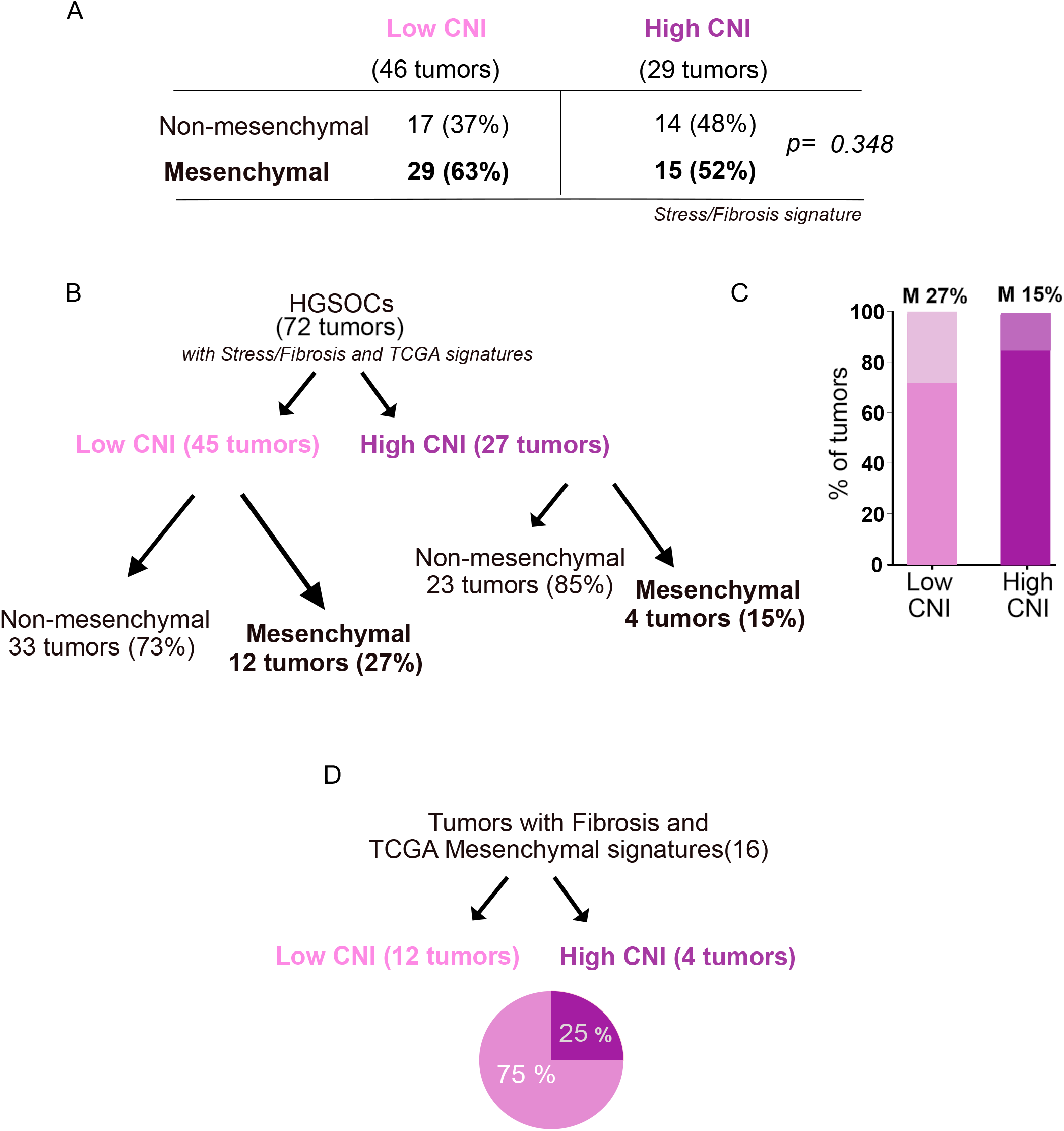
Low CNI tumors are associated with mesenchymal signatures and are of worse prognosis. (A) Contingency table summarizing the number of non-mesenchymal and mesenchymal tumors based only on Stress/Fibrosis signature (28). in the cohort used in this study according to their CNI. Statistical significance was determined by Fisher’s exact test. (B) Diagram summarizing the distribution of low and high CNI tumors and the percentage of non-mesenchymal and mesenchymal tumors in each category represented in the graph in (C). (D) Diagram representing the distribution of mesenchymal tumors in the cohort characterized in this study according to CNI.

These results show that a fraction of low CNI tumors display mesenchymal signatures. But more importantly, they also show that in the low CNI category, which is of worse prognosis according to the clinical data set presented here, non-mesenchymal signatures are also present.

## DISCUSSION

Centrosome amplification has been described as a frequent event in human tumors. Here, the analysis of a large cohort of EOCs has identified centrosome loss and an overall high frequency of low centrosome numbers as a characteristic of these tumors. Even if cells with extra centrosomes could be easily identified, their frequency was very low in the large majority of the EOCs examined. The higher frequency of low CNI tumors and the presence of cells without centrosomes has recently been described in human prostate tumors (Wang et al., 2020). Together, these two studies raise the novel possibility that at least in prostate and in ovarian cancers, centrosome loss is a frequent centrosome numerical aberration.

Interestingly, in prostate cancer, centrosome loss was associated with tumor progression, and inducing centrosome loss in non-transformed prostate epithelial cells was sufficient to generate genetic instability and malignant tumors in mice (Wang et al., 2020). In our tumor cohort, low CNI tumors did not associate with a particular tumor grade or stage or even with frequency of chromosome number alterations or structural lesions. Importantly, both the overall survival and time of relapse after chemotherapy decreased in low CNI compared to high CNI tumors. These findings, therefore, suggest a worse prognosis and outcome for patients with low CNI tumors. Significantly, our low CNI signature allows a better stratification of Homologous Recombination proficient (HRP) patients who are not sensitive to standard chemotherapy.

EOCs, and in particular HGSOCs, are highly aggressive and invasive. Patients often present tumor masses in peritoneal tissues due to cancer cell dissemination through ascites fluid (Lengyel, 2010). This process involves a particular form of invasion where EOCs clear mesothelial cells to invade through the mesentery (Iwanicki et al., 2011). Our data suggest that centrosome loss is not translated into an increased mesothelial clearance capacity or invasion through the basement membrane, at least in the experimental conditions described here. These findings suggest that yet unidentified properties of low CNI EOC cells may contribute to the observed worse prognosis.

Another relevant finding from our work relates to the enrichment of mesenchymal tumor subtype in low CNI tumors. Different molecular subgroups have been identified from transcriptomic analyses using independent cohorts of HGSOCs (Tothill et al., 2008; Bentink et al., 2012; Verhaak et al., 2012; Bell et al., 2011). Interestingly all these studies found one molecular group invariably associated with poor prognosis (Zhang et al., 2016; Kieffer et al., 2020). This group, called respectively stromal, angiogenic, or mesenchymal, was also identified in the Institut Curie cohort and referred to as “fibrosis” (Mateescu et al., 2011; Kieffer et al., 2020). The association between mesenchymal signatures and worse prognosis is partially explained by the expression of specific genes in the cancer cell population. Importantly, our study describes a new feature of cancer cells from the mesenchymal HGSOCs population-low CNI. From our analysis it is also evident that other characteristics of low CNI tumors need to be identified.

Our work paves the way for the analyses of a high number of naïve human cancers to characterize the centrosome status. It will be essential to perform this type of study in other cancers to identify the frequency of low centrosome numbers. This will help us understand whether low centrosome number is a frequent centrosome alteration among human cancers. Finally, the unexpected findings described here-the association between low CNI and poor prognosis-deserve further attention. They might represent a novel predictive biomarker that will serve to stratify patients when choosing chemotherapy regimens or in clinical assays.

## MATERIALS and METHODS

### EXPERIMENTAL MODEL AND SUBJECT DETAILS

#### Ovarian cancer cohort

All 100 ovarian cancer samples included in this study were obtained from patients treated at the Institut Curie Hospital for primitive epithelial ovarian cancer. Clinical data, including FIGO staging, were prospectively registered and summarized in Supplementary Table 1. After the pathology review of cryosections, frozen tissues were used for DNA, RNA and proteins extractions and subsequent analysis. All samples were taken before chemotherapy administration and obtained from the Biological Resource Center (BRC) of Institut Curie (certification number: 2009/33837.4; AFNOR NF S 96 900). All patients received platinum-based chemotherapy, in most cases with a combination of carboplatin and paclitaxel. Chemosensitivity was based on the classical criterion of platinum sensitivity, defined as disease progression after completion (last dose) of the platinum chemotherapy. Normal ovarian tissues were obtained from hysterectomy or prophylactic oophoro-salpingectomy.

According to French regulations, patients were informed of the studies performed on tissue specimens and did not express opposition. All analyses were approved by the National Commission for Data Processing and Liberties (N° approval: 1487390), as well as the Institutional Review Board and Ethics committee of the Institut Curie.

#### Cell culture

SKOV3 (ATCC^®^ #HTB-77) cell lines used in this study were purchased from ATCC (LGC Promochem Sarl), OVCAR8 cells were obtained from the laboratory of F. Mechta-Grigoriou. Ovarian cancer cell lines were cultured in DMEM/F12 media (ThermoFisher Scientific #31331028) supplemented with 10% Foetal Bovine Serum (FBS, Dutscher #500101L), streptomycin (100µg/mL) and penicillin (100 U/mL). The human mesothelial cell line MeT-5A was purchased from ATCC (#CRL-9444) and cultured in Medium 199 containing 1.5 g/L sodium bicarbonate (Sigma-Aldrich #M4530), 10% FBS, 3.3 nM epidermal growth factor (EGF, Sigma-Aldrich #E9644), 400 nM hydrocortisone (Sigma-Aldrich #H0888-1G), 870 nM zinc-free bovine insulin (Sigma-Aldrich #I9278), and 20 mM HEPES (Gibco #15630).

Cells were maintained at 37°C with 5% CO_2_ in the air atmosphere. They were routinely checked for mycoplasma (PlasmoTest™-Mycoplasma Detection Kit, InvivoGen, #rep-pt1) and underwent cell authentification by short tandem repeat analysis (powerplex16 HS kit, Promega #DC2101) processed at the Genomics Platform (Department of Translational Research, Institut Curie).

#### Immunofluorescence staining of centrosomes

##### Tissue sections

Frozen tissue sections of ovarian cancers and healthy tissues (20µm of thickness) were fixed in cold methanol (−20°C) for 5 min and washed 3 times for 10 minutes in PBS 1X. Sections were permeabilized 10 min using PBS supplemented with 0.5% Triton X-100, blocked 1h in PBS + 0.3% Triton X-100 + 3% of Bovine Serum Albumin (BSA). Tissues sections were incubated overnight at 4°C with primary antibodies diluted in PBS 1X + 0.3% Triton X-100 + 3% BSA. We used the PCM mouse anti-pericentrin (1/250, Abcam #ab28144) and rabbit anti-CDK5RAP2 (1/500, Bethyl #BETIHC-00063) markers. We used rabbit anti-CEP135 (1/500, generated in the lab) to recognize centrioles. Sections were washed 3 times for 10 min in PBS 1X + 0.1% Triton X-100 + 1% BSA and incubated for 6h with secondary antibodies at 4°C: goat anti-mouse IgG (H+L) highly cross-adsorbed secondary antibody Alexa Fluor 568 (1/500, Invitrogen #A-11031), goat anti-rabbit IgG (H+L) highly cross-adsorbed secondary antibody Alexa Fluor 488 (1/500, Invitrogen #A-11008). After 3×10 min of washing in PBS 1X + 0.1% Triton X-100 + 1% BSA, sections were mounted using Vectashield with DAPI mounting media (VectorLaboratories, #H-1200).

##### Cell lines

Cells were fixed in cold methanol (−20°C) for 5 min, washed and permeabilized 3 times for 5 minutes using PBS-T (PBS 1X + 0.1% Triton X-100 + 0.02% Sodium Azide). Next, cells were blocked for 30 min at RT with PBS-T supplemented with 0.5% BSA. Cells were incubated for 1h at RT with primary antibodies diluted in PBT + 0.5% BSA. We used the same antibodies as described above. Cells were washed 3 times for 5 min and incubated for 30 min with secondary antibodies diluted in PBT + 0.5% BSA: goat anti-mouse IgG (H+L) highly cross-adsorbed secondary antibody Alexa Fluor 568 (1/500, Invitrogen #A-11031), goat anti-Rabbit IgG (H+L) highly cross-adsorbed secondary antibody Alexa Fluor 488 (1/500, Invitrogen #A-11008). Cells were washed 3 x 5 min and incubated for 10 min with DAPI (1/2000, Invitrogen #D1306) diluted in PBT + 0.5% BSA. Finally, cells were washed the last three times in PBT + 0.5% BSA and once with PBS 1X, then mounted with a home-made mounting medium.

#### Stable cell lines with PLK4 inducible overexpression

##### Generation of inducible cell lines

To generate PLK4 inducible stable cell lines from OVCAR8 and SKOV3 cells, we used a doxycycline-inducible PLK4 lentiviral expression system (Holland et al., 2012). Viruses were produced in HEK293T cells, co-transfected with two other vector plasmids using lipofectamine 2000: a vesicular stomatitis virus envelope expression plasmid (Vsvg) and a second-generation packaging plasmid (pPax2). Viral particles were then used to infect OVCAR8 and SKOV3 cell lines for 24hrs. Infected cells were selected using bleomycin 50µg/mL (Santa cruz Biotechnology #sc200134A) for 15 days. Newly generated stable cell lines iOVCAR8 and iSKOV3 were then expanded in DMEM/F12 media supplemented with 10% tetracycline-free foetal bovine serum (FBS, Dutsher #S181T), streptomycin (100µg/mL,) and penicillin (100 U/mL). To induce PLK4 overexpression, cells were treated with doxycycline (1µg/mL) for 96 hrs.

##### Cell growth

10^5^ cells were plated per well in a 6-well plate and treated after adhesion with centrinone, doxycycline or corresponding controls. Living cells were trypsinized and counted at 24hrs, 48hrs, 72hrs and 96hrs post-seeding by Vi-Cell analyzer (Beckman Coulter), using a trypan blue exclusion assay.

##### Cell death

10^5^ cells were plated per well in a 6-well plate and treated with centrinone, doxycycline or corresponding controls over 96hrs. Next, cells were washed twice in cold PBS 1X and 100µl of cells suspension were stained for 15 min with 5µl of Annexin V APC and 10µl of propidium iodide 0.5mg/ml (PI), all furnished in the same kit (Biolegend #640932). Apoptotic (annexin V-positive) and necrotic (PI-positive) cells were detected using a flow cytometer (BD LSR II cytometer). FlowJo software was used to analyze results.

#### Rac1 activation assay

##### Pull-down assay

We performed Rac1–GTP pull-down assay using the Rac1 activation kit (# BK035-S, Cytoskeleton) according to the manufacturer’s instructions. After centrinone or doxycycline treatment (and corresponding controls), adherent cells were scrapped and collected in lysis buffer. Then, protein extracts were incubated with PAK-PDB affinity beads. All the experiments were done at 4 °C. Next, beads were washed and resuspended in laemmli buffer for western blotting analysis. *Western blotting:* Proteins were separated on 4-20% SDS electrophoresis gel and transferred onto PVDF membranes using Trans-Blot Turbo Transfer System (#1704156, Biorad). Images were acquired using Chemidoc Imaging system (Biorad) and band intensities were quantified using Image Lab 6.0.1 software (Biorad).

##### Transient depletion of centrosomes

For centrosome depletion, we used the centrinone drug previously described in (Wong et al., 2015) and now commercially available (Clinisciences #HY-18682). Briefly, 10^5^ iOVCAR8 or iSKOV3 cells were plated per well in 6-well plates and allowed to adhere for at least 4hrs before centrinone treatment at 200nM. Dimethyl sulfoxide (DMSO, Sigma-Aldrich #D8418) alone, at equivalent concentrations (v/v), was used as a negative control. Cells were incubated at 37°C for 96 hrs.

Note that the number of centrosomes was quantified (as described in the quantification section) following DMSO or centrinone treatment, to verify the efficiency of the drug before all functional assays.

#### Spheroid-induced mesothelial clearance assay and live cell imaging

Ovarian cancer cells were cultured for 24hrs on standard culture plates and 72hrs on Poly-2-HydroxyEthlylMethacrylate-coated culture dishes (poly-HEMA, Sigma #3932). Poly-HEMA prevents the cells from attaching to the culture dish, allowing them to remain in suspension and form spheroids. Briefly, dishes were coated with poly-HEMA at 12 mg/mL in 95% ethanol and 0.8mg/cm^2^ of density, next dried overnight at 37°C and sterilized before the experiment using ultra pure water supplemented with streptomycin (100µg/mL) and penicillin (100 U/mL). Then, poly-HEMA coated dishes were washed 3 times with ultra pure water and PBS before 10^5^ cells were plated.

To form mesothelial cells monolayer, 2.10^5^ Met-5A cells were plated in Ibidi µ-Slide 8 Well (Clinisciences #80826) coated with collagen type I (Sigma-Aldrich #C3867-1VL) and incubated for 48h at 37°C. Before imaging, Met-5A monolayer were labelled with 5 µM of CellTracker™ Orange CMRA Dye (ThermoFisher Scientific #C34551) and cancer cell spheroids were labelled with 10 µM of CellTracker™ Green CMFDA Dye (ThermoFisher Scientific #C7025) for 30min. The tumor cell spheroids were added to the mesothelial monolayer and allowed to attach for 30 min before imaging. The exclusion of mesothelial cells induced by tumor spheroids was analyzed by live imaging.

In parallel, a pool of tumor cell spheroids was dissociated using trypsin, transferred onto slides by cytocentrifugation (cytospin^TM^, Thermo Fisher Scientific) and labelled with centrosome markers (as described above) to validate treatment efficiency (centrinone *vs* DMSO) in decreasing centrosome numbers.

#### Basement membrane isolation

For animal care, we followed the European and French National Regulation for the Protection of Vertebrate Animals used for Experimental and other Scientific Purposes (Directive 2010/63; French Decree 2013-118). Mesentery BM was isolated from 5 months old female C57Bl6/N mice and glued (3M Vetbond) on 24-well plate inserts (BD Biosciences) from which the polycarbonate membrane was previously removed. Those basement membranes (BM) were decellularized for 40min into 1M ammonium-hydroxide (Sigma-Aldrich). BM are sterilized O/N at 4°C with 4ug/mL of ciprofloxacin (Panpharma) and 1.25mg/mL metronidazole (B.Braun) diluted into PBS. Mesenteries were then stored for up to 48hrs at 4°C into PBS with 2% Antibiotic-Antimycotic solution (ThermoFisher).

#### Invasion assays (BM) and staining

10^5^ iSKOV3 cancer cells were cultured for 48hrs on standard culture plates and for 48hrs on Poly-HEMA-coated 6-well plates, in presence of centrinone at 200nM or DMSO alone at an equivalent concentration (v/v).

The BM was placed into a well filled with DMEM-F12 supplemented with 10% FBS, 2% antibiotic-antimycotic, 10mM Hepes (Thermofisher). On the top side of the mesentery, aggregates from one well of a 6-well plate were plated in the presence of DMEM-F12 with 2% antibiotic-antimycotic, 10mM Hepes. Cells were cultured for 7 days at 37°C and 5% CO2 in the presence of centrinone at 200nM or DMSO at equivalent concentrations (v/v) added on both sides of the BM with each medium change after 3 days.

#### Immunofluorescence

BM was washed in PBS for 5min and fixed with 4% PFA at RT. Cells were washed in PBS three times and stained for 2hrs using Alexa Fluor™ 488 Phalloidin (1unit/mL, A12379 ThermoFisher) and DAPI (1ug/mL, D1306 ThermoFisher). BM was washed three times in PBS then mounted on glass-bottom dishes using Polymount medium (Polysciences) on both sides of the mesentery.

#### Immunofluorescence microscopy

##### For tissue sections

###### Confocal microscopy

LSM Nikon A1r was used to obtain optical sections along the Z axis (60x, Z-distance of 0.5 µm, NIS Element software) of ten random fields from the entire tissue section. Centrosomes were identified through the co-localization of two centrosomes markers.

###### Super resolution microscopy

Images were acquired on a spinning disk microscope (Gataca Systems, France), through a 100x 1.4NA Plan-Apo objective with a sCMOS camera (Prime95B, Photometrics, USA), z distance of 0.2 µm. Multi-dimensional acquisitions were performed using Metamorph 7.10.1 software (Molecular Devices, USA). Super resolution was achieved on the CSU-W1 spinning disk equipped with a super-resolution module (Live-SR, Gataca systems). Images are presented as maximum intensity projections generated with ImageJ software.

##### For cell lines

###### Fluorescence microscopy

Images were acquired on an upright widefield microscope (DM6B, Leica Systems, Germany) equipped with a motorized XY and a 100X objective (HCX PL APO 100X/1,40-0,70 Oil from Leica). For each condition, optical sections of images were acquired with a Z-distance of 0.3 µm (Metamorph software) from at least 10 random fields. Images are presented as maximum intensity projections generated with ImageJ software.

###### Live imaging

Images were acquired every 30 minutes over 12hrs using a spinning disk microscope (20x objective, 2.5 µm of z sections, Gataca Systems, France). Images were presented as maximum intensity projections generated with ImageJ software.

#### Invasion assays

Cells were imaged with an inverted laser scanning confocal LSM 880 NLO (Zeiss, Jena, Germany) coupled with Argon 488 laser (GFP) and diode 405 (DAPI) using 25x/0,8NA oil-immersion objectives (Zeiss). BM were imaged using second-harmonic generation microscopy. Optical sections of images were acquired with a Z-distance of 2.5 µm and treated using Imaris (Bitplane).

#### Analysis of genomic alterations and Homologous Recombination Deficiency (HRD)

CytoScan HD SNP-arrays (Affymetrix, ThermoFisher Scientific) data were processed using the GAP methodology to obtain absolute copy number (CN) profiles (Popova et al., 2009), including DNA structural rearrangements and chromosome number. The DNA index was calculated as the averaged CN, and tumor ploidy of tumors was set as near-diploid (DNA index <1.3) or near-tetraploid (DNA index >1.3). HRD was detected based on the number of Large-scale State Transitions (LSTs) as described previously (Popova et al., 2009). Briefly, LST was defined as a chromosomal breakpoint (change in CN or major allele counts) between adjacent regions of at least 10 Mb. The number of LSTs was calculated after smoothing and filtering out CN variant regions < 3 Mb in size based on two ploidy-specific cut-offs (15 and 20 LSTs per genome in near-diploid and near-tetraploid tumors, respectively). Tumors were classified as HRD (equal or above the cut-off) or HR Proficient (HRP) (below the cut-off). Genomic data were available for 92% (81/88 samples) of high-grade serous ovarian cancer (HGSOC).

#### Analysis of gene expression level and transcriptomic signature

mRNA expression levels were analyzed using GeneChip U133Plus 2.0 arrays (Affymetrix, ThermoFisher Scientific) as previously described in (Goundiam et al., 2015). Stress and fibrosis signatures were obtained from the laboratory of Fatima Mechta-Grigoriou, as described in (Mateescu et al., 2011). Classification of tumors from the TCGA cohort according to the DIMP signature was performed by hierarchical clustering using Euclidean distance and Ward’s agglomeration method, according to (Integrated genomic analyses of ovarian carcinoma, 2011).

#### Proliferation and mitotic indexes assessment in high-grade serous ovarian cancers

##### Proliferation index

We performed immunochemistry assays using mouse anti-human ki67 antibody (M7240, DAKO, 1/200 at pH9) in a series of paraffin-embedded tissue blocks of HGSOC. Sections of 3 µm were cut using a microtome from the paraffin-embedded tissue blocks of normal tissue and invasive lesions. Tissue sections were deparaffinized and rehydrated through a series of xylene and ethanol washes. Briefly, the key steps included: (i) antigen retrieval with ER2 pH9, (Leica: AR9640); (ii) blocking of endogenous peroxidase activity with Bond polymer refine detection kit (Leica: DS9800) (iii) incubation with primary antibodies against the targeted antigen; (iv) immunodetection with Revelation and counter staining Bond polymer refine detection kit (Leica: DS9800). Immunostaining was performed using a Leica Bond RX automated immunostaining device. We performed an immunohistochemical score (frequency x intensity) through analysis of 10 high-power fields (HPF, x 400). All quantifications were performed by 2 pathologists with blinding of patient status.

#### Mitotic index

Paraffin-embedded tissue sections of tumors were stained with hematoxylin and eosin. The mitotic count was determined by the number of mitotic figures found in 10 consecutive high-power fields (HPF), in the most mitotically active part of the tumor (entire section). Only identifiable mitotic figures were counted. Hyperchromatic, karyorrhectic, or apoptotic nuclei were excluded.

### DATA AVAILABILITY

Microarray data are available in the Gene Expression Omnibus under the accession number GSE132088.

### QUANTIFICATION AND STATISTICAL ANALYSIS

#### Centrosome quantification

##### Tumors

For each sample, 10 randomly chosen fields were considered. Using ImageJ software, we visually counted the number of nuclei and the number of centrosomes (co-localization of CDK5RAP2 and PCNT). The Centrosome Nuclei Index (CNI) was obtained by dividing the total number of centrosomes by the total number of nuclei.

##### Cell lines

The process was similar to that in the previous sections (above). At least 100 cells were quantified in different randomly chosen fields.

##### Mesothelial clearance quantification

The area-visible as a black region-induced by cancer cell spheroids invading into fluorescent mesothelial monolayer was analyzed every 30 min using ImageJ software. The area of the aperture size measured as readout of clearance was normalized by the initial spheroid size.

##### Invasion assay quantification

Analysis was performed using Imaris (BitPlane). The total number of nuclei was counted using the surface module. The BM was settled as the reference frame. Invaded nuclei were automatically counted as the object detected below the reference frame. Invasion frequency was calculated as the number of invading nuclei per the total number of nuclei per field obtained with the x25 objective.

##### Statistical analysis

All the analyses were processed using R or GraphPad Prism software. Survival analysis was performed using the Kaplan-Meier method and log-rank test for group comparison. Predictiveness curves were used to define a threshold for patient stratification according to the disease relapse at 6 months. We calculated interval confidence using a bootstrap resampling process. The Classification And Regression Trees (CART) method was used to dichotomize our population into two groups as low or high CNI tumors. A multivariable logistic regression model was used to assess the association with low or high CNI status.

For all functional analyses, we performed at least three independent experiments. Results were plotted as mean ± SEM. All data underwent normality check (Shapiro-Wilk test) and appropriate tests were performed for group comparison (Wilcoxon or Mann-Whitney; ANOVA). Fisher’s exact test was used for contingency tables to evaluate the association between parameters.

For mesothelial clearance, repeated-measure analysis of covariance (ANCOVA) was used, since longitudinal measurement of clearance were balanced with evenly space-time points for all the conditions. The clearance (mean) according to the log of time (hours) was plotted, then differences in slopes and intercepts among regression lines were evaluated. Differences were considered statistically significant at values of *P* ≤ 0.05.

## ACKNOWLEDGMENTS

We are grateful to the patients who consented to participate in this research and to the medical teams involved in their care. We thank S. Godinho, F. Gergely, S. McClelland, S. Taylor for sharing unpublished results, discussions, and/or comments on the manuscript. We thank F. Edwards, V. Marthiens, S. Gemble, A. Goupil, D. Vargas for discussions and comments on the manuscript. We thank the Tissue Imaging (PICT-IBiSA) and Nikon Imaging Centre at Institut Curie, member of the French National Research Infrastructure France-BioImaging (ANR10-INBS-04). We thank A. Vieillefon, A. Rapinat, and D. Gentien from the Genomics Platform of the translational research department at Institut Curie for cell line authentification.

The project was supported by a PIC project grant from the Labex CelTisphybio (2013, Institut Curie), INCA PL-BIO grants (2015-PLBIO15-237), WWC research grant (21-0042), Institut Curie and the CNRS. The Basto lab is a member of the CelTisphybio labex.

## AUTHOR CONTRIBUTION

The project was initially designed and conceptualized by O.G., X.S. and R.B. with significant input from S.R.R. J.P.M. and A.S. performed most experiments, including immunostaining, CNI quantifications of all the tissues and cell lines, the clearance and invasion assays; C.C was involved in setting up the protocols for centrosome analysis in cells and tissues; J.B. and C.P-G contributed to mesothelial clearance and invasion assays with significant input from DMV about dissemination processes; A.H. helped establish the stable cell lines used for functional assays; B.M. and A.L. performed predictiveness curves and provided expertise in statistical analysis. Y.K performed the analysis of TCGA DIMP transcriptomic signature. T.P. and M.H.S. helped with and advised on the HRD and ploidy status analysis. P.G. analyzed gene expression levels and provided advice in statistical analysis. G.B. and V.B. did the pathological review of all the specimens; G.B. analyzed mitotic index; D.M and A.N. analyzed ki67 in tumors; O.G. performed activation assays. O.M. managed tumor samples availability; X.S.-G. and A.V.S. provided human samples from the pathology department of Institut Curie; R.R. supplied surgical pieces to enlarge the cohort and provided expertise in ovarian cancers. F.M-G provided the OVCAR8 cell line, stress/fibrosis signature. S.R.R advice in methodology. The work was supervised by O.G and R.B.

---

**Supplementary Figure 1.**
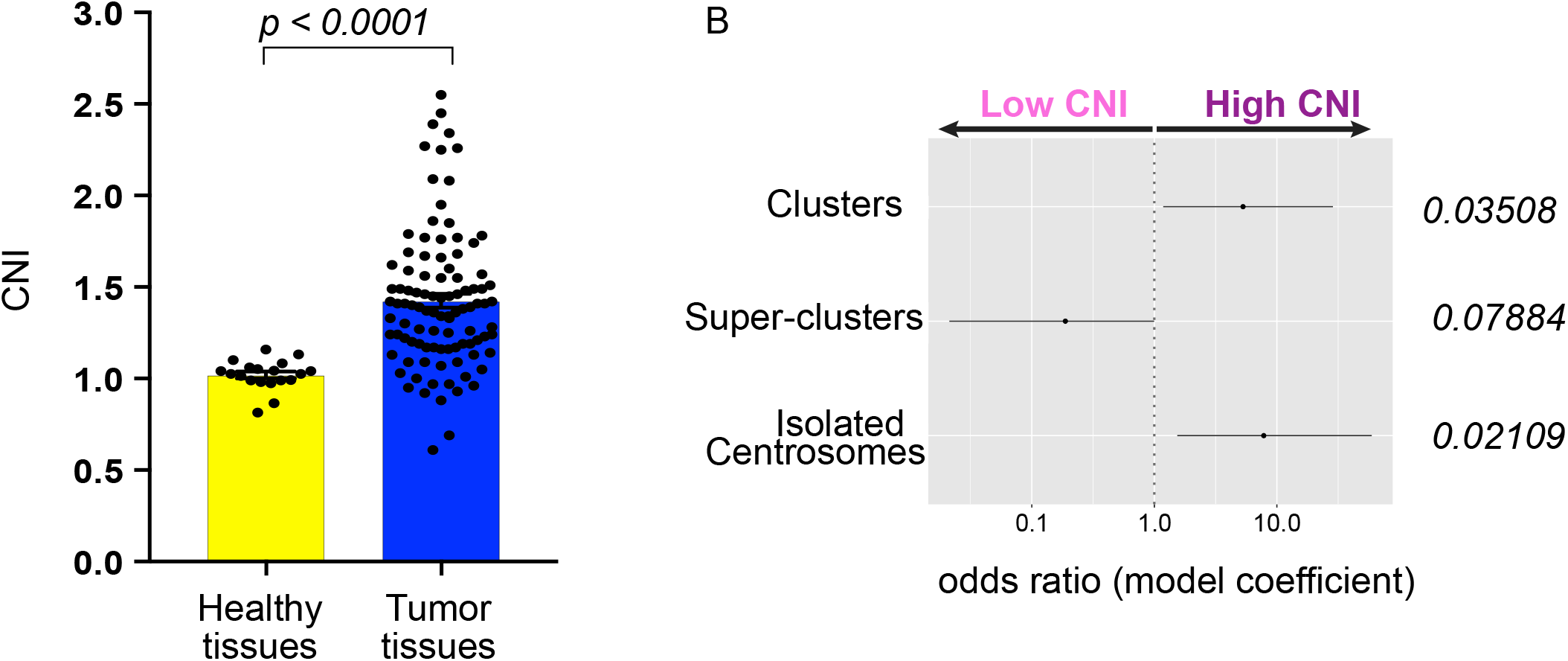
Characterization of centrosome number alterations in EOCs. (A) Graph bar of the average CNI in healthy and tumors tissues. (B) Forest plot illustrating the association of ECs categories (isolated centrosomes, clusters and super-clusters) with Low and High CNI tumors (HGSOCs), p value from multivariate logistic regression analysis.

**Supplementary Figure 2.**
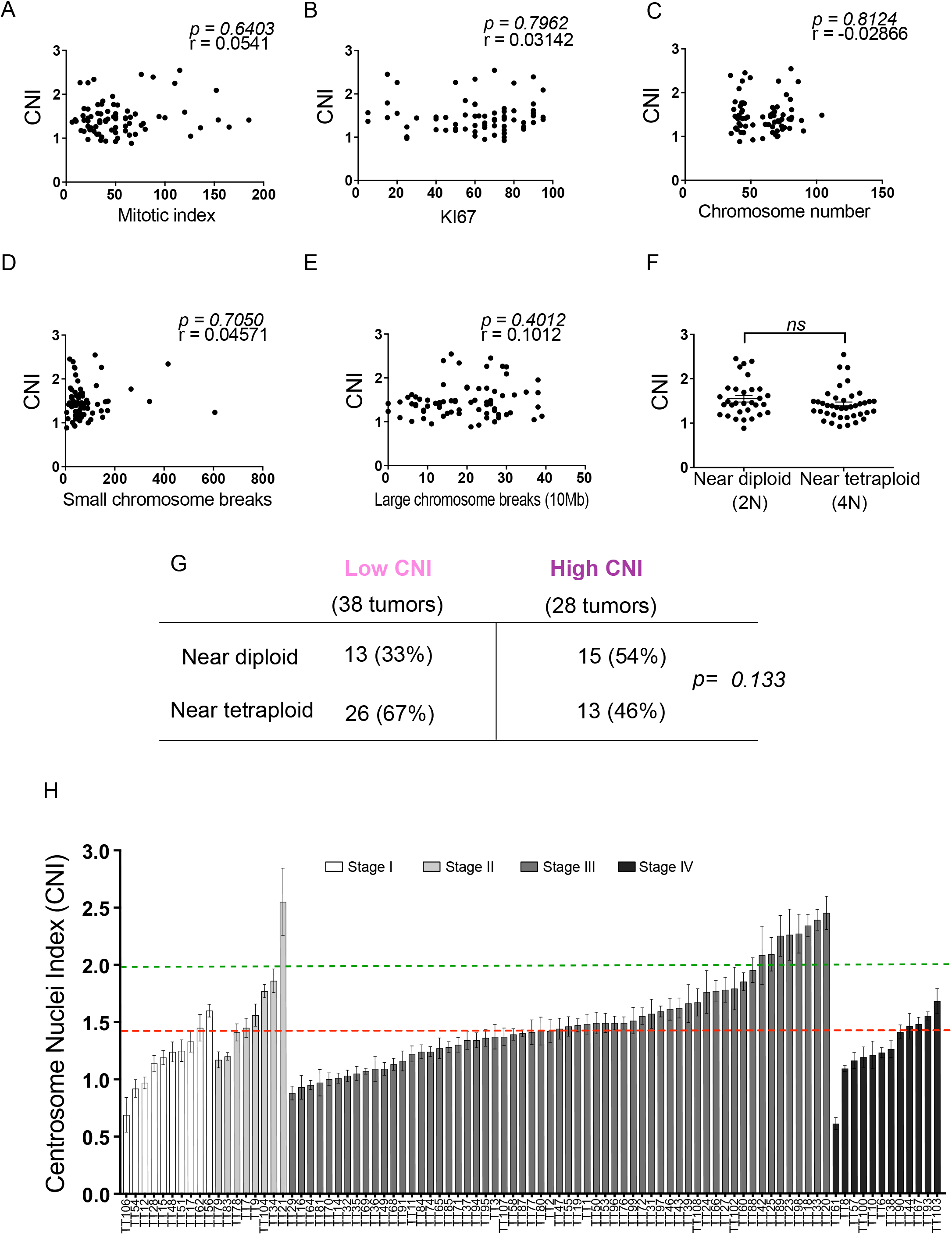
Characterization of CNI related to HGSOCs features. (A-F) Graphs showing the distribution of the mitotic index (A), Ki67 positive cells (B), chromosome numbers (C), small (D) or large (E) chromosome breaks and ploidy (F) found in HGSOCs according to the CNI, p value from Spearman correlation or Mann-Whitney tests. (G) Contingency table summarizing the association of ploidy with CNI in HGSOC tumors, Fisher’s exact test. (H) Plot showing that both low and high CNI tumors were found in all stages from I to IV, even if the cohort includes mainly grade III tumors. The red line represents the threshold of CNI value defining HGSOCs as low or high CNI tumors (1.45). The green dash line represents centrosome amplification as defined by the literature (>2 centrosomes in the cell)

**Supplementary Figure 3.**
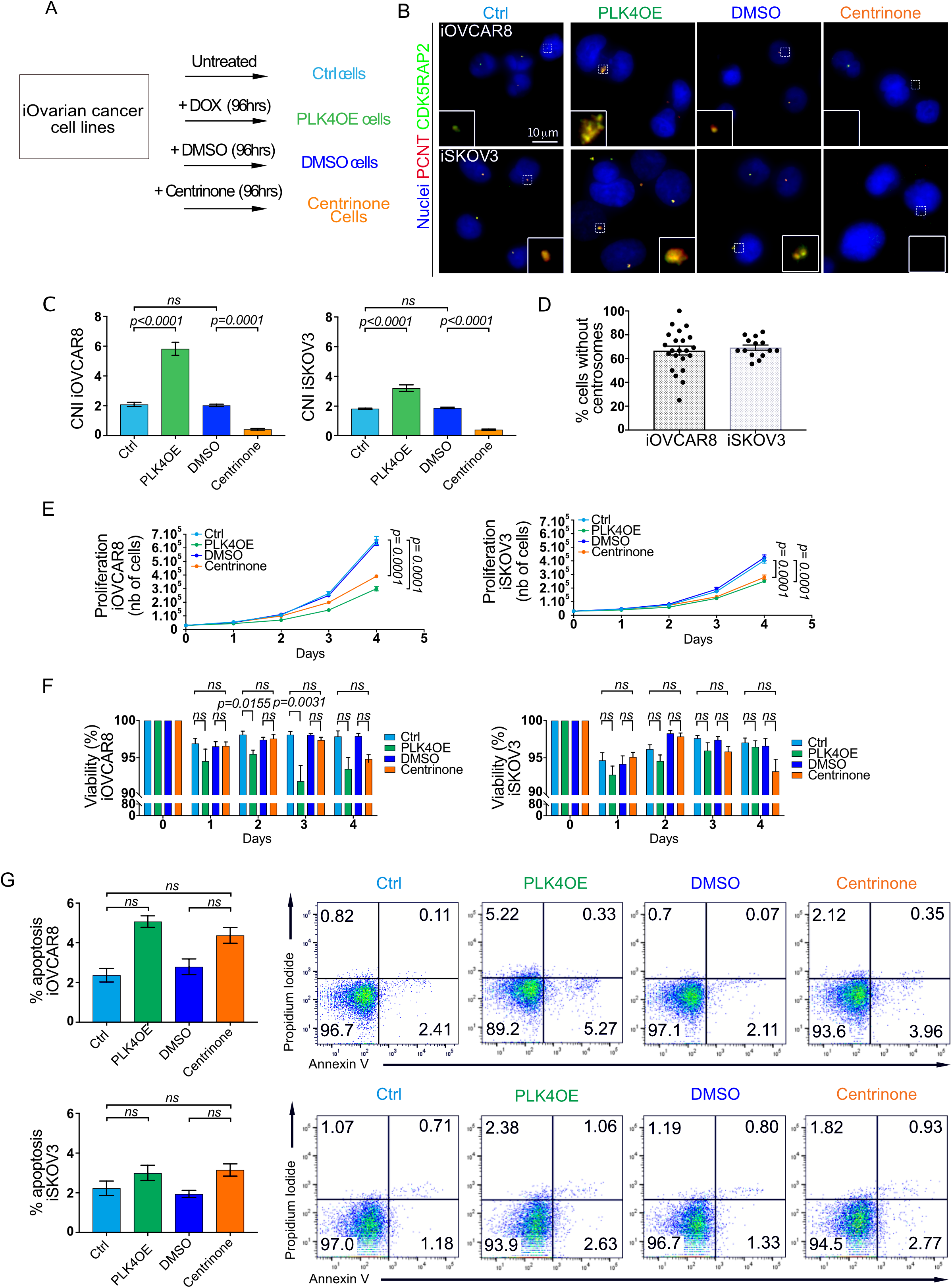
Characterization of ovarian cancer stable cell lines after gain or loss of centrosomes. (A) Scheme of the different experimental conditions and nomenclature used. (B) Representative images of iOVCAR8 (top) and iSKOV3 (bottom) after the indicated treatments labelled with antibodies against PCNT and CDK5RAP2 (red and green respectively, DNA in blue). Scale bar 10μm. The white dashed squares represent the regions shown in higher magnification on the bottom. (C) Graph bars representing the CNI after the different treatments of iOVCAR8 (left) and iSKOV3 (right). (D) Graph bar representing the quantification of the percentage of cells without centrosomes after centrinone treatment. A minimum of 150 cells were analyzed. (E) Graphs representing the proliferation of iOVCAR8 and iSKOV3 cells after each treatment for 4 days. Statistical significances assessed with two-way ANOVA. (F) Graph bars representing the viability of iOVCAR8 (top) and iSKOV3 cells after the designated treatment according to indicated timing in days. Statistical significance was assessed with one-way ANOVA. (G) On the left, graph bars represent the percentage of apoptotic cells iOVCAR8 (top) and iSKOV3 (bottom) after each indicated treatment. Statistical significances were assessed with the Wilcoxon test. On the right, representative FACS plots showing Annexin V+ (x-axis) and PI+ (y-axis) cells, for iOVCAR8 (top) and iSKOV3 (bottom) cells after the indicated treatments. For all sections, n=3 independent experiments.

**Supplementary Figure 4.**
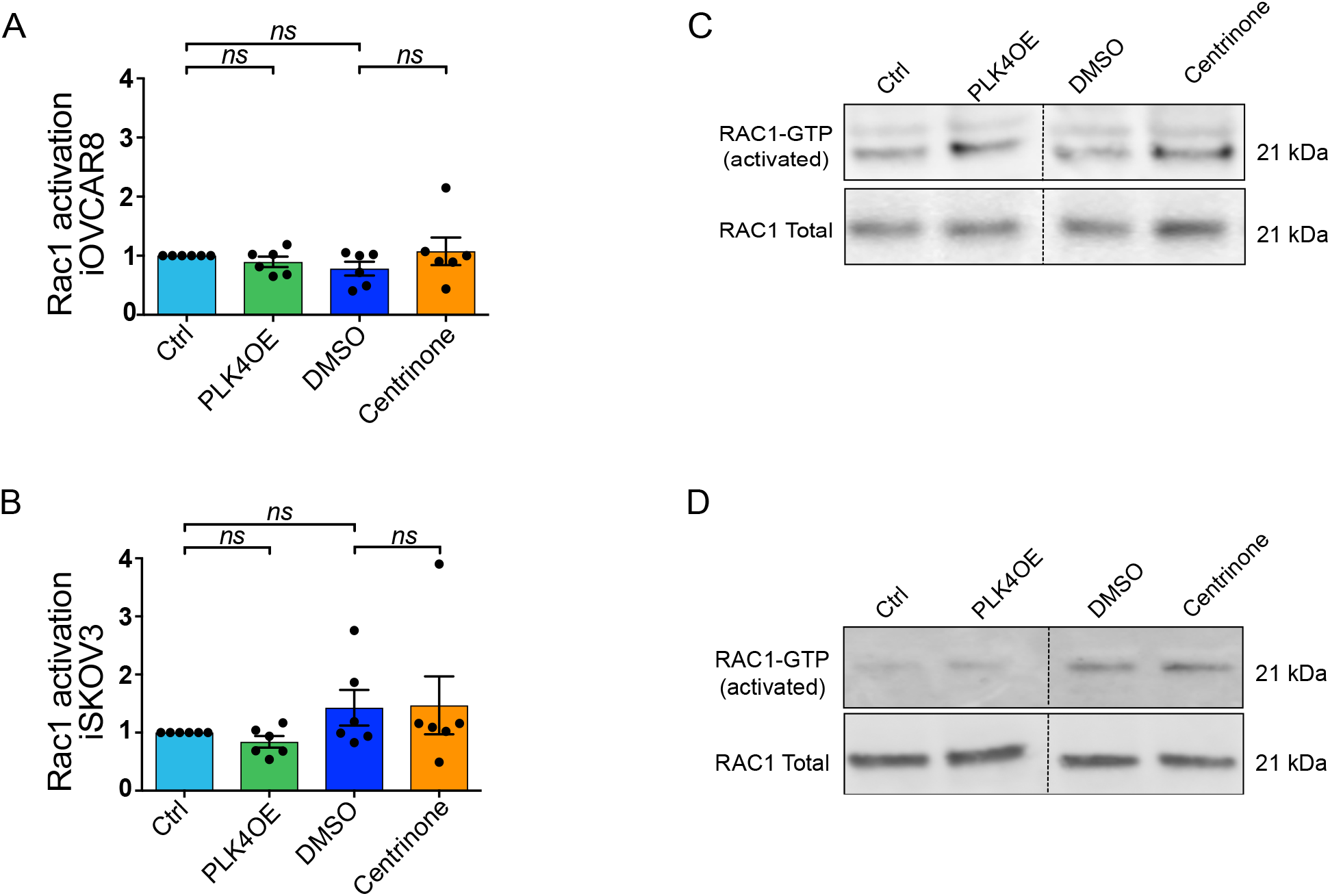
Analysis of RAC1 activation in ovarian cancer stable cell lines. (A-B) Graph bars representing the quantifications of active RAC-1 (A-B) performed by western blot (C-D) in iOVCAR8 (top) and iSKOV3 (bottom) cell lines after the indicated treatments. Statistical significances were assessed with one-way ANOVA, n=6 independent experiments.

**Supplementary Figure 5.**
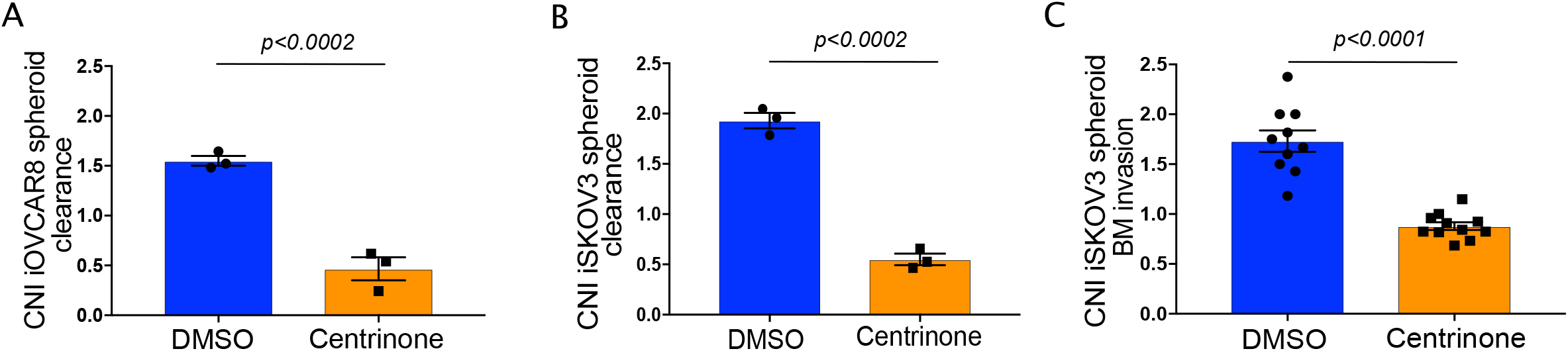
Quantification of CNI in cells used in different experiments. (A-C) Dot plot graphs showing the CNI quantifications in (A-B) mesothelial cell clearance and (C) BM assays. Statistical methods were determined using t-tests.

**Supplementary Table 1.**
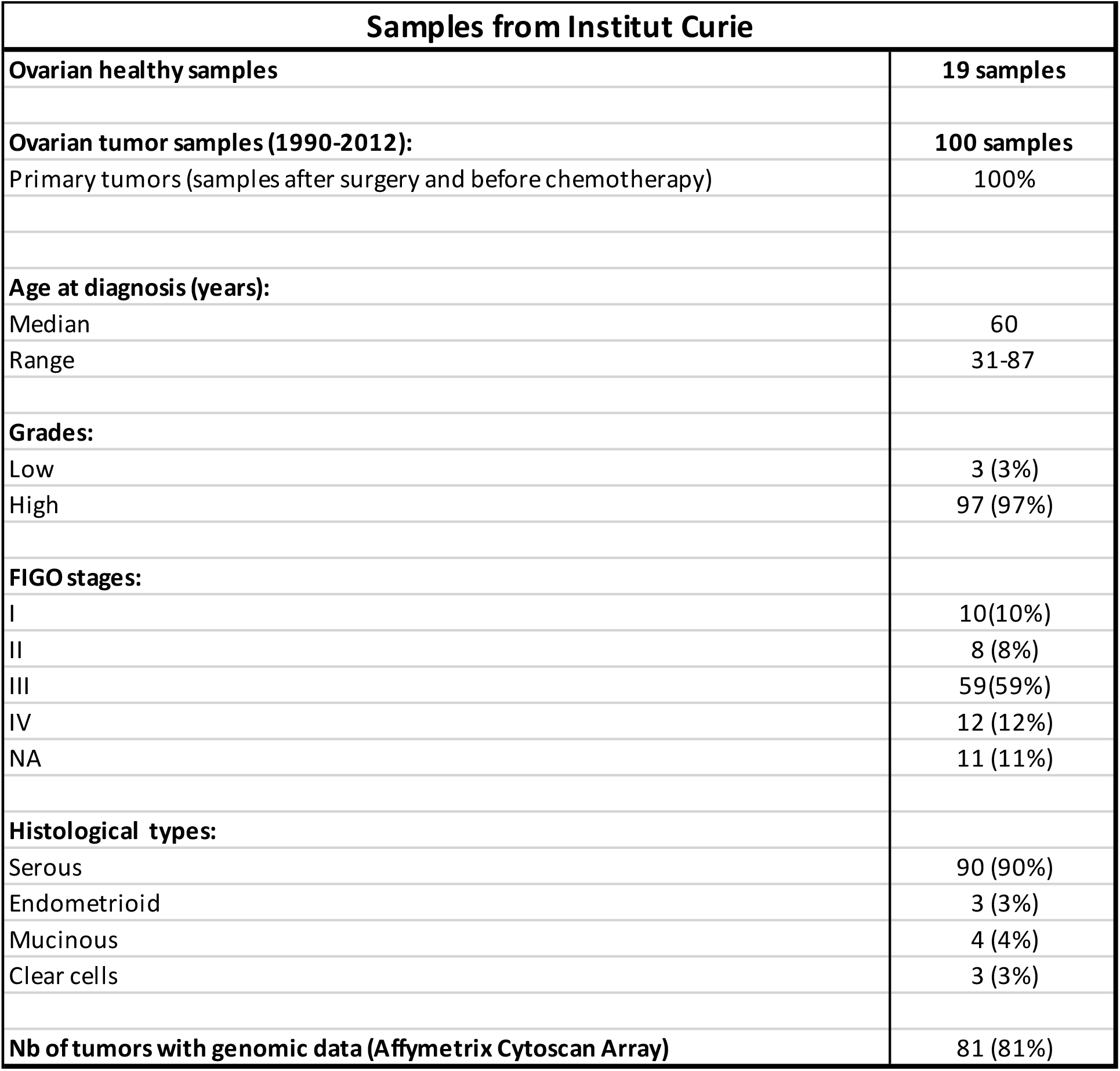

| Samples from Institut Curie |  |
| --- | --- |
| **Ovarian healthy samples** | **19 samples** |
| **Ovarian tumor samples (1990-2012):** | **100 samples** |
| Primary tumors (samples after surgery and before chemotherapy) | 100% |
| **Age at diagnosis (years):** |  |
| Median | 60 |
| Range | 31-87 |
| **Grades:** |  |
| Low | 3 (3%) |
| High | 97 (97%) |
| **FIGO stages:** |  |
| I | 10(10%) |
| II | 8 (8%) |
| III | 59(59%) |
| IV | 12 (12%) |
| NA | 11 (11%) |
| **Histological types:** |  |
| Serous | 90 (90%) |
| Endometrioid | 3 (3%) |
| Mucinous | 4 (4%) |
| Clear cells | 3 (3%) |
| **Nb of tumors with genomic data (Affymetrix Cytoscan Array)** | **81 (81%)** |

